# A resistome roadmap: from the human body to pristine environments

**DOI:** 10.1101/2021.10.08.463752

**Authors:** Lucia Maestre-Carballa, Vicente Navarro-López, Manuel Martínez-Garcia

## Abstract

A comprehensive characterization of the human body resistome (sets of antibiotic resistance genes (ARGs)) is yet to be done and paramount for addressing the antibiotic microbial resistance threat. Here, we study the resistome of 771 samples from five major body parts (skin, nares, vagina, gut and oral cavity) of healthy subjects from the Human Microbiome Project and addressed the potential dispersion of ARGs in pristine environments. A total of 28,731 ARGs belonging to 344 different ARG types were found in the HMP proteome dataset (n=9.1×10^7^ proteins analyzed). Our study reveals a distinct resistome profile (ARG type and abundance) between body sites and high inter-individual variability. Nares had the highest ARG load (≈5.4 genes/genome) followed by the oral cavity, while the gut showed one of the highest ARG richness (shared with nares) but the lowest abundance (≈1.3 genes/genome). Fluroquinolone resistance genes were the most abundant in the human body, followed by macrolide-lincosamide-streptogramin (MLS) or tetracycline. Most of the ARGs belonged to common bacterial commensals and multidrug resistance trait was predominant in the nares and vagina. Our data also provide hope, since the spread of common ARG from the human body to pristine environments (n=271 samples; 77 Gb of sequencing data and 2.1×10^8^ proteins analyzed) thus far remains very unlikely (only one case found in an autochthonous bacterium from a pristine environment). These findings broaden our understanding of ARG in the context of the human microbiome and the One-Health Initiative of WHO uniting human host-microbes and environments as a whole.

**Importance:** The current antibiotic resistance crisis affects our health and wealth at a global scale and by 2050 predictions estimate 10 million deaths attributed to antibiotic resistance worldwide. Remarkably, a comprehensive analysis of ARG diversity and prevalence in different human body sites is yet to be done. Undoubtedly, our body and human built-environment have antibiotic resistant bacteria than can also be transported to other environments. Hence, the analysis of Human Microbiome Project dataset provides us not only the opportunity to explore in detail the ARGs diversity and prevalence in different parts of our body but also to provide some insights into the dispersion of ARGs from human to natural populations inhabiting pristine environments. Thus, our data would help to stablish a baseline in ARG surveillance protocols to asses further changes in antibiotic resistances in our society.

## Introduction

Since the discovery of antibiotics, human and animal health has profoundly changed. Undoubtedly, antibiotics have not only saved millions of lives but also have transformed modern medicine (Ventola, 2015; Centers for Disease Control and Prevention, 2019). Nevertheless, antibiotic overuse, incorrect prescription, extensive use of antibiotics in agriculture and farming and the low availability of new antibiotics have led to a major antibiotic resistance crisis, wherein bacterial pathogens are becoming resistant to available antibiotics (Ventola, 2015). In the US, it has been estimated that antibiotic-resistant organisms cause 2.8 million infections and 35,900 deaths each year (Centers for Disease Control and Prevention, 2019). This not only is a health issue but also affects food security and requires significant financial investment. For instance, it has been estimated that in 2017, the annual treatment of six multidrug-resistant bacteria costs approximately $4.6 billion to the US healthcare system (Nelson *et al*., 2021). By 2050, predictions estimate that over 10 million deaths and a total cost of ≈100 trillion USD will be attributed to antibiotic resistance worldwide (Brogan and Mossialos, 2016; O’Neill, 2016), and recently, the WHO estimated that in 10 years, antimicrobial resistance could force up to 24 million people into extreme poverty (IACG, 2019).

Antibiotic resistance is a natural process in which bacteria become resistant to antibiotics using different mechanisms, which are classified as phenotypic resistance (due to physiological changes and nonhereditary) or acquired (when antibiotic resistance is genetically gained) (Olivares *et al*., 2013). Different antibiotic resistance genes (ARGs) that confer resistance to antibiotics could be acquired due to mutations in the bacterial genome or through horizontal gene transfer (HGT). The transference of ARGs could be mediated by bacteria, viruses, plasmids or even vesicles (Emamalipour *et al*., 2020). Among the possible antibiotic classifications, the most common are the ones based on their chemical structure (drug classes, e.g, tetracycline, beta-lactams, aminoglycosides…), mode of action (determined by the antibiotic target, mainly proteins, cell membrane and nucleic acids) and spectrum of activity (from narrow to broad) (Wright, 2010; Etebu and Arikekpar, 2016; Reygaert, 2018).

The occurrence of antibiotic resistance has increased and accelerated since antibiotics are constantly present in the environment, derived from anthropogenic sources such as wastewater treatment plants, hospitals or domestic use (Rodriguez-Mozaz *et al*., 2020). Another cause of this increase is the dispersion of resistant bacteria from hot spots, such as wastewater treatment plants (from our own human microbiome) and built environments (i.e., microorganisms found in human-constructed environments) (Baron *et al*., 2018; Centers for Disease Control and Prevention, 2019), which continuously disseminates our microbes and thus parts of their genetic content.

The human microbiome project (HMP) (Methé *et al*., 2012), an interdisciplinary effort, was developed with the objective of characterizing the human microbiome. For this purpose, samples from different major body parts of healthy humans were obtained and sequenced, producing one of the largest resources for the study of the human microbiome (Huttenhower *et al*., 2012). To the best of our knowledge, comprehensive analysis and cross-comparison of the human resistome from all human body parts studied within the HMP have not been performed, but to date, some valuable but separate and non-interconnected studies have been performed for the oral cavity and the skin (Carr *et al*., 2020; Li *et al*., 2021). Addressing the abundance and diversity of ARGs as a whole in all human body parts has enormous potential to broaden our knowledge on the dispersion of ARGs from human bacteria within different microbial populations in nature.

Thus, here, in the context of antibiotic resistance-related health concerns, in addition to analyzing in detail the antibiotic resistance genes present in the HMP-studied body sites, we studied the potential spread of ARGs from the human body to different types of pristine environments. These environments are supposed to be undisturbed and not affected by anthropic actions. While many places, such as caves or polar environments with no apparent and visible human activity, are often perceived as pristine environments, human activity unfortunately leaves an indirect ever-increasing footprint. Here, we use some of these pristine environments as a model to address and estimate the potential “mobility” of common human ARGs found in the human body to better assess the global impact of antibiotic resistance in our ecosystems, in line with the One Health concept (i.e., human health and animal health are interdependent and bound to ecosystems) (Atlas, 2012). Pristine environments are commonly used as “reporter ecosystems” to monitor pollution and climate change and, in our case, specifically to measure how deep the potential impact of the spread of antibiotic resistance is.

## Results

### Comprehensive metagenomic characterization of the human resistome

The HMP (Huttenhower *et al*., 2012) aimed to characterize the diversity and metabolic potential of the microbiomes of healthy human subjects from different body sites. In this study, we analyzed the resistome (i.e., pool of antibiotic resistance genes) of these body parts, examining a total of 771 HMP samples from the oral cavity, skin, nares, vagina and gut (Suppl. Table 1 and 2). Detection of ARGs was performed using amino acid sequence similarity searching against the following reference ARG databases (see methods for details): CARD 3.0.3 (Jia *et al*., 2017), RESFAMS (Gibson *et al*., 2015) and ARG_ANNOT (Gupta *et al*., 2014). ARGs with an e-value ≤ 10^−5^, amino acid identity ≥ 90% and bit score ≥ 70 were considered *bona fide* ARG hits.

From all the detected HMP proteins (9.17E+07), a total of 28,731 ARG hits were found, representing between 0.02 and 0.08% of the relative abundance of HMP proteins depending on the body site analyzed (Suppl. Table 2). Overall, nearly all analyzed samples (99%) from the different human body sites showed at least one ARG. The exceptions were some specific HMP samples from the nares (≈14% of nares samples), skin (4.25% of skin samples) and vagina (45% of vagina samples), in which no ARG was detected (Suppl. Table 2).

On average, tetracycline resistance genes were the most abundant antibiotic resistance genes in the HMP dataset (Fig. 1A and B), followed by MLS or fluoroquinolone resistance genes, the ranks of which were dependent on the analyzed body site. Tetracycline resistance genes were the most abundant ARGs in vagina (53.4%) and gut (40.52%), whereas their abundance decreased in oral cavity, skin and nares (26.02, 9.03 and 10.45%), where the most dominant antibiotic resistance genes were, respectively, fluoroquinolone (30%), multidrug (18.22%) and beta-lactamase resistance genes (≈20%) (Fig. 1A). In gut samples, the second most abundant resistance genes were the ones conferring resistance against beta-lactamases (as in skin), while in the vagina, the second most abundant were multidrug resistance genes (19.37%). Aminocoumarin resistance genes were only found in the gut, while peptide antibiotic resistance genes were found in all body parts analyzed and they were more frequent in skin and nares (representing a 6-fold increase compared with the relative abundance of this antibiotic class resistance gene in the rest of the body).

**Fig. 1.**
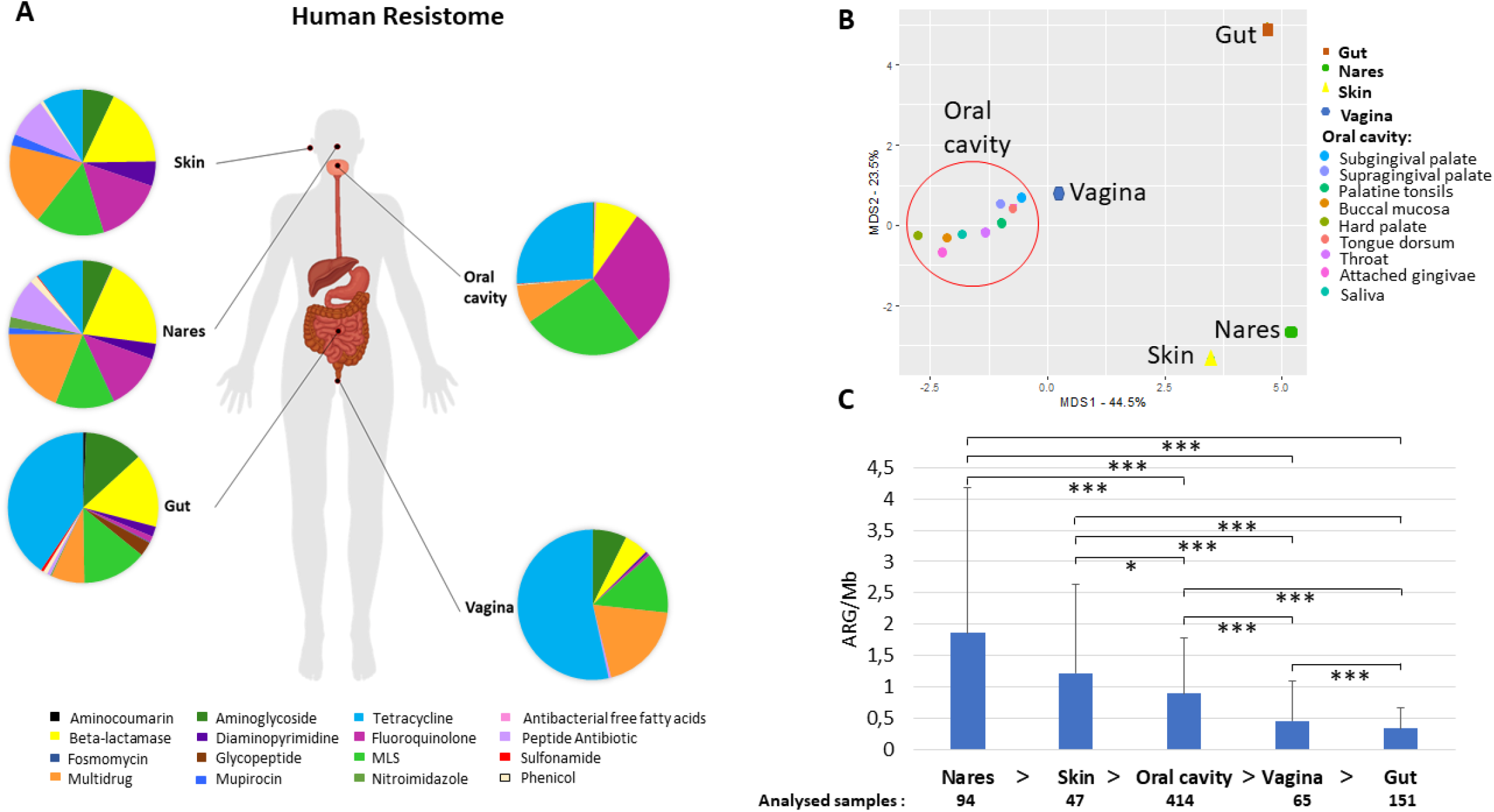
Human Resistome. Human atlas of the ARGs grouped by drug class their confer resistance to present in different body parts. The body groups studied were the gut, the skin (retroauricular crease), vagina (posterior-fornix, mid vagina and vagina intraotus), the nares and the oral cavity (hard palate, buccal mucosa, saliva, subgingival plaque, attached gingivae, tongue dorsum, throat, palatine tonsils, and supragingival plaque) (A). PCoA analysis of the different body sites distributed according to their relative abundance of AR to different drug classes (B). The samples included in the group oral cavity (hard palate, buccal mucosa, saliva, subgingival plaque, attached gingivae, tongue dorsum, throat, palatine tonsils, and supragingival plaque -shaped as a circle-) gathered together and separately from nares, skin and gut samples. Abundance of antibiotic resistance genes calculated as ARGs hits per assembled Mb and number of samples included in each body group (C). Welch test was performed to compare ARG abundances between different body sites. All paired samples showed statistically significant differences but the nares and the skin. P-values (P) considerer as significant were indicated with an asterisk: P ≤ 0.05 *, P ≤ 0.01**, P ≤ 0.001***.

All samples from the oral cavity (hard palate, buccal mucosa, saliva, subgingival and supragingival plaque, attached/keratinized gingivae, tongue dorsum, throat and palatine tonsils) showed a similar pattern of resistance to the different antibiotic classes with minor variations (Fig. 1B; Suppl. Fig. 1). Separated from the oral cavity, the skin and nares showed similar dominant antibiotic resistance genes grouped by drug classes (fluoroquinolone, multidrug, macrolide-lincosamide-streptogramin (MLS) and beta-lactamase resistance), although the ARG in nares displayed resistance to 14 different drug classes, while ARG present in skin displayed resistance to 10 different drug classes. The bacteria from the vagina had resistance against 8 antibiotic classes, being the lowest number of the 5 body parts compared in this study (the top three ARG ranked were tetracycline, fluoroquinolone and MLS resistance genes). Remarkably, nares and gut showed resistance to the highest number of antibiotic classes (14 out of the 16 different classes found in this study).

As shown in Fig. 1C, the body part that had the highest abundance of ARGs per assembled mega-base pair (Mb) was the nares (1.86±2.32 ARGs/Mb), followed by the skin (1.22±1.42 ARGs/Mb) and oral cavity (0.90±0.88 ARGs/Mb) (Fig. 1C). It is worth noting that the gut (0.34±0.33 ARGs/Mb) had the lowest amount of ARGs per Mb among all the analyzed body parts (Fig. 1C). The Welch test employed to compare the abundance of different body parts showed statistically significant differences (P-value≤0.05) between almost all body parts but not between the skin and nares (Fig. 1C). No significant differences were found between male and female subjects in any of the body sites analyzed (Suppl. Fig. 2). According to recent estimates of the average genome size (AGS) of human microbes from different body parts of HMP datasets (Nayfach and Pollard, 2015), in general, the correlation of ARGs and the AGS indicated that the number of ARGs per bacterial genome ranged from 1.3 in stool (AGS=3.9 Mb) to 3 in nares (AGS≈2.5 Mb).

### Characterization of dominant antibiotic resistance genes in the human body

In this section, beyond the above-described diversity and abundance of antibiotic resistance gene classes in the HMP dataset, we sought to study in detail the pool of different types of ARGs and the identity of antibiotic-resistant microbes harboring these ARGs. Within each of the antibiotic classes, different types of ARGs are described in databases (2404 in CARD (Jia *et al*., 2017), 2038 in ARG_ANNOT (Gupta *et al*., 2014), and 3169 in RESFAMS (Gibson *et al*., 2015)). In addition, based on the antibiotic mechanism of action (Reygaert, 2018), five types of antibiotics are defined: antibiotics that 1) inhibit cell wall synthesis (e.g., beta-lactams), 2) depolarize the cell membrane (e.g., lipopeptides), 3) target nucleic acid synthesis (e.g., quinolones), 4) inhibit metabolic pathways (e.g., sulfonamides) and 5) affect protein synthesis (e.g., MLS antibiotics or tetracyclines) (Reygaert, 2018). Therefore, ARGs could provide protection against one specific antibiotic or different types of antibiotics.

In the HMP datasets, after comparison with all three of the above ARG databases, a total of 344 different type of ARGs were found in all the analyzed samples (Suppl. Table 3 and 4). The gut samples had 198 different ARGs, the highest number and diversity among the analyzed body sites, while the lowest ARG diversity was found in the vagina (46) (Suppl. Table 4). The most abundant type of ARG in the oral cavity was *patB*, which provides resistance to fluoroquinolones via antibiotic efflux (ARO: 3000025). The *fmtC* gene was the predominant ARG in the nares and skin, while *tetQ* was the most common ARG in the gut and in the vagina, the most frequent gene was tetM (Suppl. Table 4).

Regarding the identification of the most common antibiotic resistant (AR) bacteria in HMP datasets (Fig. 2) based on the best-hit score, as expected, the results differed among body parts. The oral cavity had 326 different species harboring ARGs, followed by the gut (257 different species). The skin showed the lowest number of different species with ARGs (a total of 52) (Fig. 2). *Streptococcus mitis* was the most abundant AR bacterium in the oral cavity. In the gut, the most abundant AR bacterium was *Escherichia coli*, while in nares and skin, *Staphylococcus* was the predominant AR bacterium (*Staphylococcus aureus* in nares and *Staphylococcus epidermis* in skin). Finally, *Gardnerella vaginalis* was the most abundant resistant species in the vagina. *S. aureus, E. coli* and *Bacteroides fragilis* were the most abundant AR bacteria found in all body sites (Fig. 2). Remarkably, from the total ARG hits found in the HMP (n=28,731), only one example detected in the oral cavity was detected with high confidence in a viral genome fragment of a human herpesvirus carrying *APH(4)-Ia*; this ARG was an aminoglycoside phosphotransferase that inactivates aminoglycosides (human subject 765560005, buccal mucosa, Suppl. Table 5).

**Fig. 2.**
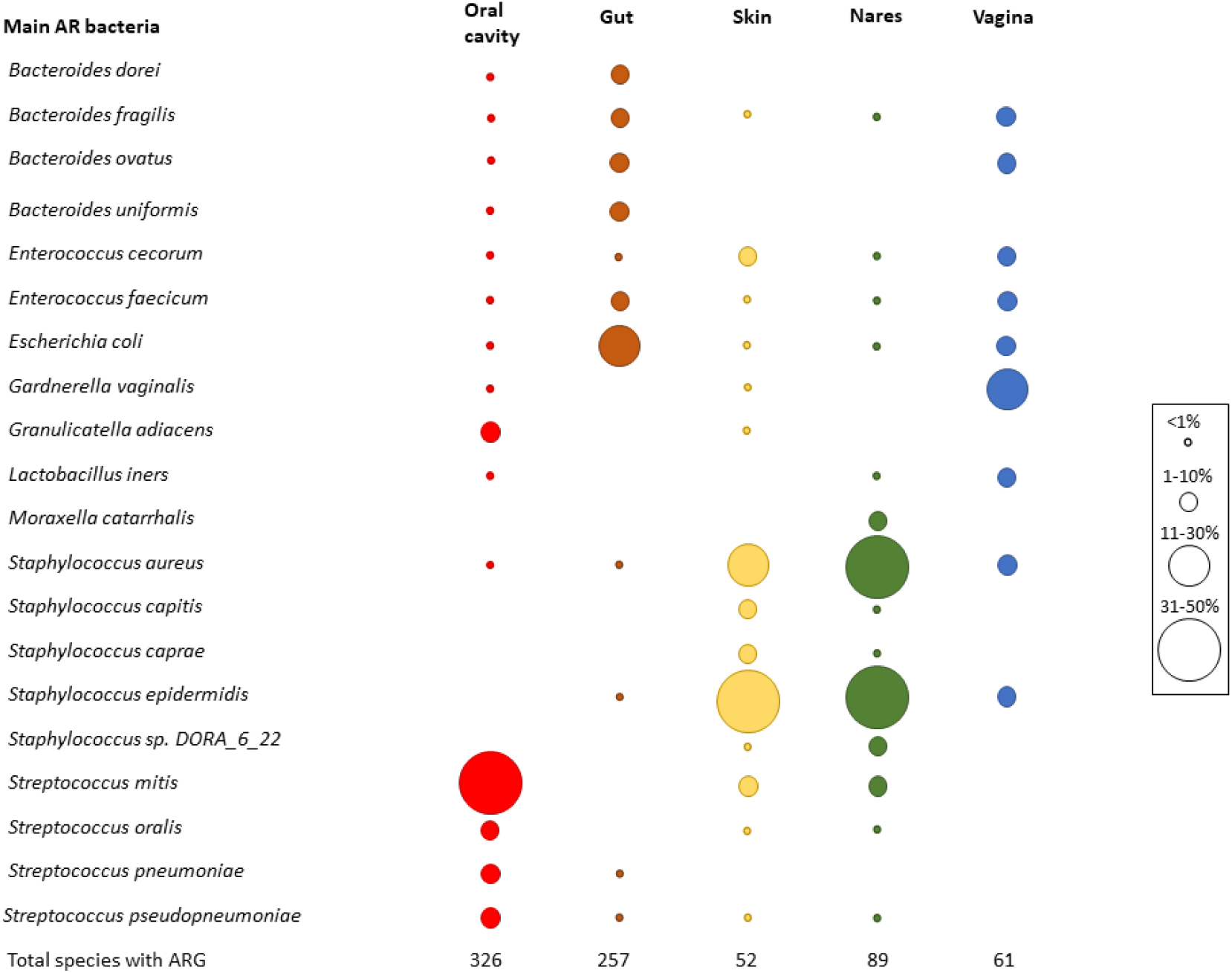
Main antibiotic resistant bacteria in HMP dataset. Relative abundance of the most abundant resistant bacteria. Top five bacteria were chosen in each body part and then the graphic was completed with the relative frequency of all the chosen bacteria in all body parts. Circle sizes were different to determine the relative abundance of each species and colours were used to differentiate the body parts (red-oral cavity, brown-gut, skin-yellow,green-nares,blue-vagina). At the bottom of the graphic the number of different species that carried ARGs is shown.

The increasing multiantibiotic-resistant bacteria (MRBs) are a major threat to human health. We next attempted to identify genome fragments (i.e. contigs) having two or more ARGs which provide insights into multiple-antibiotic resistance (MR) bacteria in HMP datasets. For this purpose, we applied the criteria for the detection of more than one ARG in the same assembled genome fragment, conferring resistance to at least two different antibiotics classes in each of the analyzed HMP samples. The percentage of metagenomic samples from the HMP with the presence of MR was between ≈25% (oral cavity) and 6% (vagina). Twenty-one percent of the analyzed gut samples had >1 contig conferring multi-antibiotic resistance potential, whereas in the skin, the percentage was 19%, and in nares, 15% of the samples showed MR (Fig. 3A). The MR frequency changed depending on the studied group. Vagina samples showed the highest multi-antibiotic resistance-related contig frequency, with a large difference among vaginal samples (0.42±0.27 MRB/assembled Mb). The skin, oral cavity and gut had the lowest frequency of MR (Fig. 3B).

**Fig. 3.**
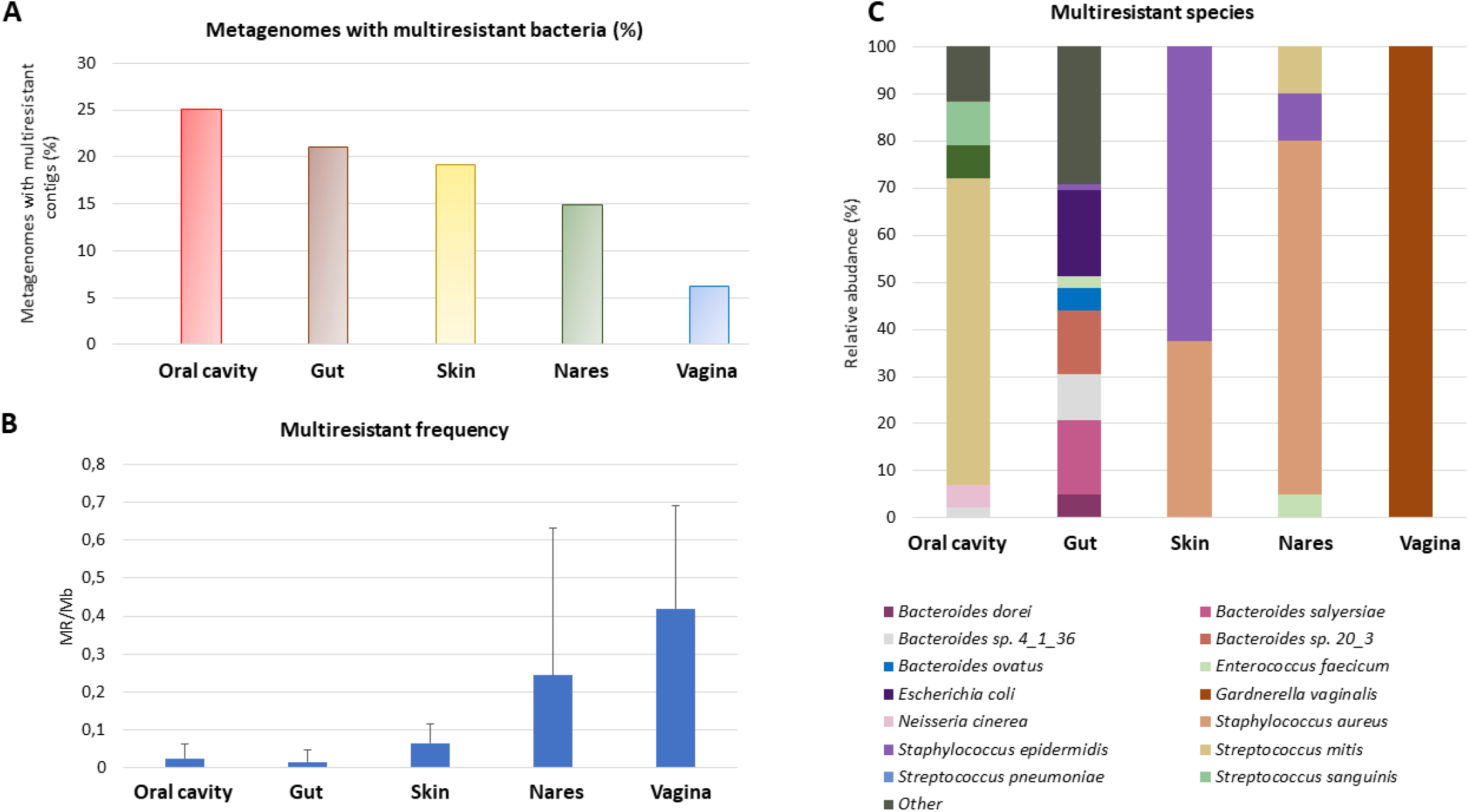
Multiresistance in the human body. Those assembled genome fragments (i.e. contigs) that had more than one ARG conferring resistance to at least 2 different antibiotic families were considered as multiresistant (MR). Percentage of metagenomes with multiresistant contigs compared with all the metagenomes studied from the same HMP sample (A). Study of the multiresistant contigs frequency in metagenomes with at least one multiresistant contig (B), to compare the different samples, the number of multiresistant contigs was divided by the assembled Mb. Standard deviation is shown in the graphic. Relative abundance of the most abundant MR (C). Only MR whose relative abundance was, at least in one body site, equal or greater than 5% were represented.

The most abundant MR species in each body site were the same in all cases and were also detected as the most predominant resistant bacteria (Fig. 3C). The MR profile was different depending on the sampling site. In the vagina there was only one MR species whereas in skin, there were only 2 main species carrying more than one ARG, while the gut had 23 MR species, with the highest number of different MR found in all body sites. None of the MRB species were found in all the body parts. In fact, 6 species (*Bacteroides* sp. *4_1_36, Bacteroides fragilis, Enterococcus faecium, Staphylococcus aureus, Staphylococcus epidermidis, Streptococcus mitis*) out of the 33 MRBs found were in two or three different parts of the body, while the rest were only body site specific (Suppl. Fig. 3).

When all of the above ARGs detected in healthy humans were clustered (≥90% amino acid identity) to study a highly conserved core of shared ARGs, it was observed that there were 3 common ARGs in all the body parts. One, MFS-type efflux protein (*msrD*), was related to resistance to macrolides. The other 2 genes were related to ribosomal resistance against tetracycline (*tetO* and *tetQ*) associated with conjugative plasmids or transposons. In feces, *tetO* was found not only in bacteria belonging to the phylum Firmicutes (*Clostridiales bacterium VE202-13;* Ga0104838_1543581) but also in bacteria of the phylum Actinobacteria (*Trueperella pyogenes MS249;* Ga0111491_10662371).

### Detection of Human Microbiome Project ARGs in pristine environments

Resistomes and ARGs dispersion from hot spots such as wastewater plants or hospitals to downstream aquatic environments have been extensively studied and characterized (Rowe *et al*., 2017; Ju *et al*., 2018; Khan *et al*., 2019). Although the presence of ARGs in environments with low or scarce human intervention has been explored (Van Goethem *et al*., 2018; Naidoo *et al*., 2020), to the best of our knowledge, it has never been explored whether common ARGs from HMP datasets are present in autochthonous bacteria from different pristine environments.

To determine the presence of ARGs from the HMP in pristine environments (Fig. 4; polar, desert, cave, hot spring, and submarine volcano environments; Suppl. Table 6) with no *a priori* anthropogenic influence, proteins from 271 different pristine environments (i.e. metagenomic datasets) were screened to search for ARGs detected in the analyzed HMP samples. Only those proteins with identity ≥90%, with bit-score ≥ 70 and belonging to genomic scaffolds with at least 4 proteins were considered for further analysis. It is important to remark that if an ARG from the HMP dataset is detected in a genome fragment from a pristine environment, two different hypothesis could be considered: this detected ARG in pristine environments was 1) an allochthon HMP gene dispersed from anthropic areas that was acquired by autochthonous bacteria inhabiting the pristine environment, or 2) this ARG in pristine environments is actually the result of contamination during sample manipulation, collection or post-processing (e.g., DNA contaminant fragments in reagents from kits, DNA sequencing and other metagenomics-related experiments) and thus is not truly present in these pristine environments.

**Fig. 4.**
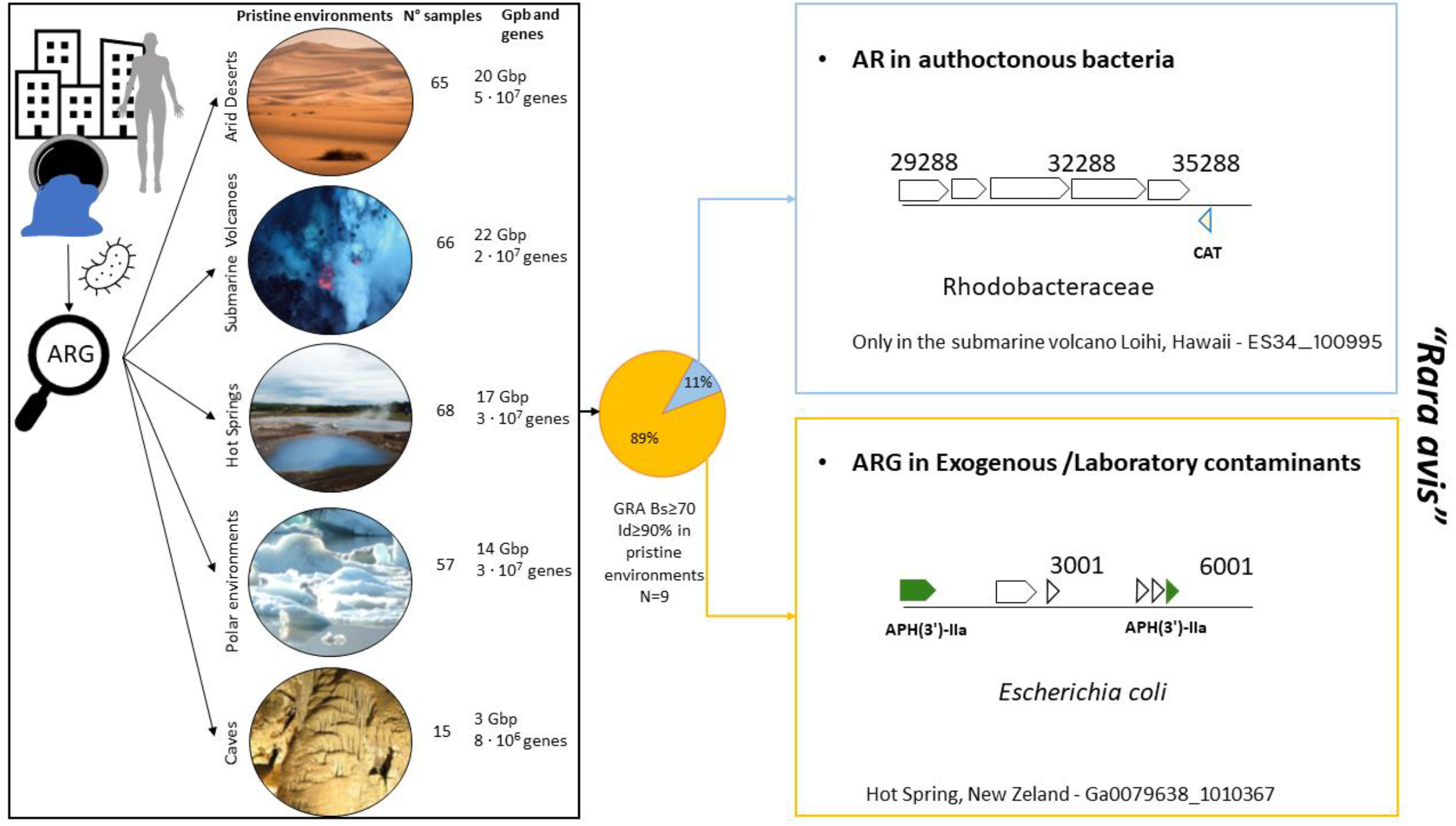
Detection of ARGs from Human Microbiome Project dataset in pristine environments (arid deserts (n=65), submarine volcanoes (n=66), hot springs (n=68), polar environment (n=57) and caves (n=15)). Only 9 ARGs were found in pristine environments according to our criteria (see methods and results). The only case of ARG found in an autochthonous bacterium in pristine environments was that of a chloramphenicol acetyltransferase (CAT) gene belonging to *Salmonella* sp. (100% identity with Ga0111015_155701; a nares sample) present in a marine bacterium found in Loihi (a submarine volcano) from the family *Rhodobacteraceae*. The presence of CAT from *Enterobacteriaceae* in *Rhodobacter* has been previously described in the coastal water of Jiaozhou Bay, (Dang *et al*., 2008). Chloramphenicol-resistant bacteria often harbor plasmids carrying the CAT gene (Shaw *et al*., 1979) that could have been transferred to *Rhodobacter*. Desert photo taken from Boris Ulzibat (PEXELS). Submarine volcano photograph courtesy of NOAA / NSF / WHOI page (https://oceanexplorer.noaa.gov/facts/volcanoes.html).

In the 271 analyzed samples from pristine environments (a total of 77 Gb of sequencing information and 2.1E+08 analyzed proteins), we detected a total of 9 ARGs from HMP dataset. Only one of those ARGs were found in a genome fragment of a putative autochthonous bacterium from the family *Rhodobacteraceae* recovered in a submarine volcano (Fig. 4; Suppl. Table 7; a chloramphenicol acetyltransferase gene 100% amino acid identical with the HMP gene dataset). The rest and great majority of detected ARGs in pristine environments were simply exogenous contaminant present in these metagenomes from manipulation or laboratory reagents. For instance, it is obvious that *Escherichia coli* should not be detected in hot springs. However, we indeed found ARGs in *E. coli* genome fragments in the corresponding hot spring metagenomes (Fig. 4, bottom right panel).

## Discussion

The human resistome has received increased attention in recent years due to its impact in our society. Usually, resistome studies focus their attention on one body site, usually studying the gut (Hu *et al*., 2013; Palleja *et al*., 2018) or, more recently, the skin (Li *et al*., 2021). To our knowledge, only one study has compared resistome traits from the gut with different parts of the oral cavity, examined via different protocols (Carr *et al*., 2020). In our study, the advantage of using only HMP samples that were subjected to standardized procedures was the elimination of biases and variability introduced by contrasting procedures from different surveys (Huttenhower *et al*., 2012). Here, in our study, we found that the most abundant ARGs in the HMP resistome provided resistance against fluoroquinolone, MLS and tetracycline resistance genes, followed by multidrug resistance genes and beta-lactamases. Members of these antibiotic classes were among the most commonly prescribed oral antibiotics in 2010 (Hicks *et al*., 2013), right before the samples were obtained, which shows a plausible relation between the consumed antibiotics and the detected resistance in American subjects, even though we cannot rule out the influence of antibiotics consumed through the food (Salyers *et al*., 2004). A human gut study from Chinese, Spanish and Danish subjects showed that more than 75% of the ARGs were tetracycline resistance genes, MLS resistance genes and beta-lactamases (Hu *et al*., 2013). This was consistent with our data since these three antibiotic classes accounted for 61% of the relative abundance found in our study with HMP samples (Van Boeckel *et al*., 2014). The characterization of resistomes from metagenomic data can also be performed from unassembled data (Arango-Argoty *et al*., 2018; Maestre-Carballa *et al*., 2019). Here, the analysis from unassembled data (Suppl. Fig. 5; Suppl. Table 8) showed that major ARGs grouped by drug classes relative abundance was similar to the one obtained with assembled data shown in Fig. 1. Even though, we cannot rule out that the normalized abundances (no. of ARG per Mb) could be biased by the metagenomic assembly step or by the very short lengths of Illumina DNA reads and the predicted amino acid sequences obtained from the HMP datasets (Suppl. Fig. 6).

The different physiological conditions, bacteria-host interactions, and average genome size (AGS) (Nayfach and Pollard, 2015) present in each body part could be important factors contributing to the differences in ARG abundance, which were statistically significant for different paired body parts analyzed, except in the skin-nares pair (Fig. 1C). In addition, the well-known inter-individual variability in the human microbiome was also observed here for ARG abundance. The highest ARG abundance was found in the nares, a body entrance for microorganisms carried by air, which could include pathogenic bacteria such as *Legionella* or *Mycobacterium* species. Airborne bacteria could also carry ARGs (Li *et al*., 2018); therefore, antibiotic resistance genes could first arrive at the nares. It has been calculated that we inhale 7 m^3^ of air and 10^4^-10^6^ bacterial cells per cubic meter of air per day (Kumpitsch *et al*., 2019) albeit the quantity varies depending on different factors, such as geography, weather, micro-niches and air pollution (Li *et al*., 2018; Kumpitsch *et al*., 2019; Zhang *et al*., 2019). In addition, seasonal variation in bacterial species in the nares environment has been observed (Camarinha-Silva *et al*., 2012). However, considering that bacteria present on the nares surface, in contrast to gut or oral bacteria, are not typically in “direct contact” with antibiotics, it is certainly surprising that the nares microbiome maintains the highest rate of ARG abundance, and more intriguing are the mechanisms used to acquire and fix these ARGs.

As shown in this study, the numbers of ARGs per assembled Mb in the gut was lower than that in the other body parts, but the ARG richness was greater. This observation is consistent with the results of Carr et al. (2020), who compared oral and fecal samples. Even though the abundance was measured with other parameters (reads per kilobase of read per million (RPKM) and coverage greater than 90%), the ARG abundance in stool was smaller than that in oral samples from China, the USA and Fiji but not western Europe (Carr *et al*., 2020). Carr et al. (2020) hypothesized that different niches in the oral cavity, such as the dorsum of the tongue, could aid the deposition of debris and microbes or even the formation of biofilms, which are structures that favor HGT between different species (Giaouris *et al*., 2015).

Regarding the bacterial species with antibiotic resistance, consistent with other studies, we found that commensals such as *Staphylococcus aureus* and *Staphylococcus epidermidis* in skin were the top 10 AR bacteria (Li *et al*., 2021). Further, some of them, such as *S. aureus*, were multidrug-resistant bacteria, with a total of 4 ARGs in nares (*arlS, arlR, dha-2, mprF*) conferring resistance to 3 different antibiotic classes (fluoroquinolone, beta-lactam and cationic antibiotics). Additionally, as expected, methicillin-resistant *S. aureus* (MRSA), which is listed among the CDC’s Antibiotic Resistance Threats in the United States (Centers for Disease Control and Prevention, 2019), was present naturally in nares from different subjects. Another species found in oral HMP samples listed in the AR Threats report was *Streptococcus pneumoniae*.

Remarkably, our data give hope, since the dispersion of ARGs detected in the HMP dataset to pristine environments is extremely infrequent and anecdotical, with only one ARG in an autochthonous bacterium among dozens of millions of analyzed genes. Therefore, even using more relaxed thresholds, it can be considered as a *rara avis* event. As shown in Fig. 4, nearly all detected ARGs from pristine environments actually belonged to laboratory contaminants or exogenous bacteria that were not obviously found in these habitats (e.g., *E. coli* in hot springs). Sometimes, a general metagenomic analysis could mislead the interpretation of the data if sequencing and genomic assembled data is not carefully inspected. Our study exemplifies very well this bias since an initial ARG search detection indeed discovered a certain number of ARGs, but later on, it was demonstrated that they were clearly not naturally present in these pristine environments.

Finally, it is important to discuss potential caveats and biases of our study. Here, we have used sequence similarity-based searches with strict conservative thresholds for detecting ARGs in metagenomics datasets to avoid false positives. Only hits with amino acid identity ≥90% and bit-score ≥ 70 against ARGs deposited in curated reference antibiotic resistance databases were considered. This methodology has been widely used in previous publications (Van Goethem *et al*., 2018; Chng *et al*., 2020; Lira *et al*., 2020). Obviously, unknown ARGs yet to be discovered and therefore not present in reference ARG databases cannot be detected using our methodology. Likewise, probably our approach has ruled out some actual ARGs present in samples that display a score similarity below our thresholds (i.e. false negatives). However, it is worth-noting, as highlighted in previous studies using our methodologies, that more rigorous thresholds are clearly preferred. It is very interesting to read the discussion on how using less strict thresholds when detecting ARGs in viruses can profoundly mislead data interpretation (Enault *et al*., 2017)). This is even more important when analyzing datasets from pristine environments since a conservative approach is preferable over using riskier thresholds. Even though we have used strict thresholds to detect only *bona fide* ARGs, it may be noted that some genes could “scape” this filter. For instance, some housekeeping genes (constitutive genes required for basic cellular functions) only require one or few mutations to conferring antibiotic resistance (e.g. *rplS, gyrA, parY*). For instance, the mutant version of the housekeeping gene *gyrA* found in common antibiotic resistance databases used in this study, typically display a very short motif called “QRDR” that is responsible for quinolone resistance (Avalos *et al*., 2015; Jia *et al*., 2017). However, in our search in HMP datasets, despite having high similarity and above our thresholds, the great majority of detected *gyrA* proteins in HMP did not have this motif (Suppl. Fig. 4) and therefore was totally unclear whether they confer antibiotic resistance. Similar cases were found for other housekeeping genes, even when they displayed high sequence similarity. Thus, to avoid including false positives that would overestimate ARG abundance, housekeeping gene hits were ruled out from our analysis.

Overall, our study provides a comprehensive analysis of the human microbiome resistomes from different body sites studied by the HMP consortium, providing valuable biological insights that can serve as baseline for further studies and be thus integrated into AMR surveillance protocols to determine the fate of the diversity and abundance of ARGs in the long term. Our data also show that the level and impact of ARGs spreading and selection pressure to fix these alleles in non-anthropogenic areas is negligible. However, it is in our hands, as a society, to control these selection pressures and, if possible, reverse and ameliorate the impact of ARGs in nature.

## Experimental Procedures

### Sample collection

A total of 751 shotgun-sequenced samples from 15 different parts of the body from healthy American adults belonging to the Human Microbiome Project (HMP) (Huttenhower *et al*., 2012) were retrieved from JGI-IMG/ER (Chen *et al*., 2021) (Suppl. Table 1). Not all HMP assembled data present at JGI-IMG/ER was accessible, thus only the available metagenomes were included in this study. The data were organized in 5 groups: Skin (retro-auricular crease), Nares, Gut, Vagina (posterior-fornix, mid vagina and vagina introitus), and oral cavity (hard palate, buccal mucosa, saliva, subgingival plaque, attached gingivae, tongue dorsum, throat, palatine tonsils, and supragingival plaque).

20 metagenomes belonging to left and right retro-auricular crease that could not be found in JGI-IMG were downloaded from the HMP page (https://www.hmpdacc.org/HMASM/) and included with the rest of HMP samples (Suppl. Table 1).

Proteins of 271 metagenomes from pristine environments (or environments with no or little human presence) were downloaded from JGI-IMG, and they were organized in 5 groups: Arid deserts (65), submarine volcanoes (66), hot springs (68), polar environments (57) and caves (15) yielding a total of 76 Gb (Suppl. Table 6). Environments associated to a host (e.g., tubeworms) were also discarded.

### HMP resistome in silico analysis

Proteins from 751 samples of the HMP were retrieved from de JGI-IMG/ER (Chen *et al*., 2021). In addition, 20 assembled metagenomes were downloaded directly from the HMP official page since they were not available in JGI. ORF of the genomic sequences downloaded from HMP were predicted with prodigal 2.6.3 (Hyatt *et al*., 2010).

Then, all obtained proteins were compared using BLASTp 2.8.1+ with the following antibiotic resistance protein databases: CARD 3.0.3 (Jia et al., 2017), ARG-ANNOT (Gupta *et al*., 2014) and RESFAMS (Gibson *et al*., 2015). Aiming to identify only high-confidence ARG, only those ARG with e-value ≤ 10^−5^, amino acid identity ≥ 90% and bit-score ≥ 70 with the mentioned ARG databases were considered as hits, thus, being more conservative than other accepted thresholds (bit-score ≥ 70) (Enault *et al*., 2017).

The ARG were grouped, following CARD 3.0.3 annotation (Jia *et al*., 2017), according to the drug class their confer resistance to. The taxonomic affiliation was extracted from the annotation found in JGI/IMG-ER (Chen *et al*., 2021). To compare the obtained results, hits were normalized by the assembled Megabase pair (Mb).

Multiresistant contigs were manually curated, and only those with at least 2 different ARG conferring resistance to at least two different drug classes were included in the analysis. The abundance of metagenomes with multiresistant contigs was calculated by dividing the number of metagenomes with at least one multiresistant contig by the total number of metagenomes studied. The frequency of multiresistant species only in metagenomes with more than one multiresistant bacteria was done by dividing the number of multiresistant by the total number of contigs.

The presence of common ARGs in all the analysed parts of the body was performed using CD-HIT (90% identity) (Fu *et al*., 2012) which was used to cluster all the ARGs found in the HMP.

For studying the effect of our threshold in housekeeping genes that requires few mutations to become resistant, gyrA proteins from the HMP dataset that were considered as ARG by our analysis were extracted and aligned against the gyrA fluroquinolone resistant gene deposited in CARD (Jia *et al*., 2017) from *Mycobacterium tuberculosis* (>gb|CCP42728.1|+|Mycobacterium tuberculosis gyrA conferring resistance to fluoroquinolones [Mycobacterium tuberculosis H37Rv]) and gyrA^R^ obtained from RESFAMS (Gibson *et al*., 2015) (NC_002952_2859949_p01) from *Staphylococcus*. The alignment was performed with MUSCLE available in Geneious 9.1.3.

### HMP antibiotic resistance genes in pristine environments

To determine the presence of ARG from the HMP in pristine environments with presumptive low or none human presence, the ARGs obtained from the human samples were compared with the proteins from the chosen metagenomes using BLASTp 2.8.1+. Only those hits with an amino acid identity ≥ 90%, a bit-score ≥ 70 and e-value ≤ 10^−5^ were considered. The taxonomic annotation was retrieved from JGI-IMG/ER only for those scaffolds with at least 4 proteins to ensure the detection of HMP ARGs in autochthonous bacteria (3 out of, at least, 4 proteins should be from the autochthonous bacteria). All the hits were manually curated to avoid false positives, especially those produced by housekeeping genes. Those belonging to taxons that could not be associated with a specific environment were discarded.

Alignments were performed using the software geneious 9.1.3.

Comparison between ARGs present in assembled and raw data was performed analysing paired unassembled and assembled metagenomes from 5 gut samples from different subjects (Subjects ID: 159005010, 159247771, 159369152, 763961826 and 246515023; Suppl. Table 7) and from five buccal mucosa samples from 5 different subjects (Subjects ID: 370425937, 764325968, 604812005, 246515023 and 809635352; Suppl. Table 7). ARG in assembled data were detected with blastp as mentioned above. ARGs detection in raw data was performed with two different strategies: DeepARG (Arango-Argoty *et al*., 2018), a machine learning algorithm that detects ARGs and normalises it by the number of 16S rRNA gene (90% identity, e-value ≤ 10^−10^), and comparing the reads with blastx against the antibiotic resistance databases CARD (Jia *et al*., 2017), ARG-ANNOT (Gupta *et al*., 2014) and RESFAMS (Gibson *et al*., 2015) (e-value ≤ 10^−5^, amino acid identity ≥ 90% and bit-score ≥ 70) and normalised by the unassembled Mb.

### Statistical analysis

One-way ANOVA was performed to compare the ARG abundance (ARG/Mb) in each body site between samples from women and men.

Comparison between ARGs hits/Mb was performed with Welch test and pairwise.t.test in R (R Core Team, 2014). P-value≤0.05 was considered as significant in all the statistical test performed.

PcoA analysis was performed calculating the distance matrix using the Euclidean distance and plotted with ggplot (Wickham, 2016). For the different sites of the body it was studied the relative abundance of each ARG categorized by antibiotic class resistance.

## Acknowledgements

We thank the support of Hospital Elche Crevillente Salud SL to conduct this research. We thank Yoshiko Misumi and Springer Nature Author Services team for their English edition.

## Funding

This project was supported by the Office of the Vice President for Research, Development and Innovation (University of Alicante) with a grant from the program “predoctoral training in collaboration with companies” (ref. UAIND18-05). Funds were also provided by Hospital Elche Crevillente Salud SL (ref. HOSPITALECLHE1-18Y).

## References

Arango-Argoty, G., Garner, E., Pruden, A., Heath, L.S., Vikesland, P., and Zhang, L. (2018) DeepARG: a deep learning approach for predicting antibiotic resistance genes from metagenomic data. Microbiome 6: 23.

Atlas, R.M. (2012) One Health: Its Origins and Future. 1–13.

Avalos, E., Catanzaro, D., Catanzaro, A., Ganiats, T., Brodine, S., Alcaraz, J., and Rodwell, T. (2015) Frequency and Geographic Distribution of gyrA and gyrB Mutations Associated with Fluoroquinolone Resistance in Clinical Mycobacterium Tuberculosis Isolates: A Systematic Review. PLoS One 10: e0120470.

Baron, S.A., Diene, S.M., and Rolain, J.M. (2018) Human microbiomes and antibiotic resistance. Hum Microbiome J 10: 43–52.

Van Boeckel, T.P., Gandra, S., Ashok, A., Caudron, Q., Grenfell, B.T., Levin, S.A., and Laxminarayan, R. (2014) Global antibiotic consumption 2000 to 2010: An analysis of national pharmaceutical sales data. Lancet Infect Dis 14: 742–750.

Brogan, D.M. and Mossialos, E. (2016) A critical analysis of the review on antimicrobial resistance report and the infectious disease financing facility. Global Health 12: 8.

Camarinha-Silva, A., Jáuregui, R., Pieper, D.H., and Wos-Oxley, M.L. (2012) The temporal dynamics of bacterial communities across human anterior nares. Environ Microbiol Rep 4: 126–132.

Carr, V.R., Witherden, E.A., Lee, S., Shoaie, S., Mullany, P., Proctor, G.B., et al. (2020) Abundance and diversity of resistomes differ between healthy human oral cavities and gut. Nat Commun 11: 1–10.

Centers for Disease Control and Prevention (2019) Antibiotic resistance threats in the United States. Centers Dis Control Prev 1–113.

Chen, I.M.A., Chu, K., Palaniappan, K., Ratner, A., Huang, J., Huntemann, M., et al. (2021) The IMG/M data management and analysis system v.6.0: New tools and advanced capabilities. Nucleic Acids Res 49: D751–D763.

Chng, K.R., Li, C., Bertrand, D., Ng, A.H.Q., Kwah, J.S., Low, H.M., et al. (2020) Cartography of opportunistic pathogens and antibiotic resistance genes in a tertiary hospital environment. Nat Med 2020 266 26: 941–951.

Dang, H., Ren, J., Song, L., Sun, S., and An, L. (2008) Dominant chloramphenicol-resistant bacteria and resistance genes in coastal marine waters of Jiaozhou Bay, China. World J Microbiol Biotechnol 24: 209–217.

Emamalipour, M., Seidi, K., Zununi Vahed, S., Jahanban-Esfahlan, A., Jaymand, M., Majdi, H., et al. (2020) Horizontal Gene Transfer: From Evolutionary Flexibility to Disease Progression. Front Cell Dev Biol 8: 229.

Enault, F., Briet, A., Bouteille, L., Roux, S., Sullivan, M.B., and Petit, M.-A. (2017) Phages rarely encode antibiotic resistance genes: a cautionary tale for virome analyses. ISME J 11: 237–247.

Etebu, E. and Arikekpar, I. (2016) Antibiotics: Classification and mechanisms of action with emphasis on molecular perspectives. Int J Appl Microbiol Biotechnol Res 4: 90–101.

Fu, L., Niu, B., Zhu, Z., Wu, S., and Li, W. (2012) CD-HIT: Accelerated for clustering the next-generation sequencing data. Bioinformatics 28: 3150–3152.

Giaouris, E., Heir, E., Desvaux, M., Hébraud, M., Møretrø, T., Langsrud, S., et al. (2015) Intra- and inter-species interactions within biofilms of important foodborne bacterial pathogens. Front Microbiol 6: 841.

Gibson, M.K., Forsberg, K.J., and Dantas, G. (2015) Improved annotation of antibiotic resistance determinants reveals microbial resistomes cluster by ecology. ISME J 9: 207–16.

Van Goethem, M.W., Pierneef, R., Bezuidt, O.K.I., Van De Peer, Y., Cowan, D.A., and Makhalanyane, T.P. (2018) A reservoir of “historical” antibiotic resistance genes in remote pristine Antarctic soils. Microbiome 6: 40.

Gupta, S.K., Padmanabhan, B.R., Diene, S.M., Lopez-Rojas, R., Kempf, M., Landraud, L., and Rolain, J.-M. (2014) ARG-ANNOT, a new bioinformatic tool to discover antibiotic resistance genes in bacterial genomes. Antimicrob Agents Chemother 58: 212–220.

Hicks, L.A., Taylor, T.H., and Hunkler, R.J. (2013) U.S. Outpatient Antibiotic Prescribing, 2010. N Engl J Med 368: 1461–1462.

Hu, Y., Yang, X., Qin, J., Lu, N., Cheng, G., Wu, N., et al. (2013) Metagenome-wide analysis of antibiotic resistance genes in a large cohort of human gut microbiota. Nat Commun 4: 1–7.

Huttenhower, C., Gevers, D., Knight, R., Abubucker, S., Badger, J.H., Chinwalla, A.T., et al. (2012) Structure, function and diversity of the healthy human microbiome. Nature 486: 207–214.

Hyatt, D., Chen, G.-L., LoCascio, P.F., Land, M.L., Larimer, F.W., and Hauser, L.J. (2010) Prodigal: prokaryotic gene recognition and translation initiation site identification. BMC Bioinformatics 11: 119.

IACG (2019) No time to wait: securing the future from drug-resistant infections report to the secretary-general of the united nations.

Jia, B., Raphenya, A.R., Alcock, B., Waglechner, N., Guo, P., Tsang, K.K., et al. (2017) CARD 2017: expansion and model-centric curation of the comprehensive antibiotic resistance database. Nucleic Acids Res 45: D566–D573.

Ju, F., Beck, K., Yin, X., Maccagnan, A., McArdell, C.S., Singer, H.P., et al. (2018) Wastewater treatment plant resistomes are shaped by bacterial composition, genetic exchange, and upregulated expression in the effluent microbiomes. ISME J 1.

Khan, F.A., Söderquist, B., and Jass, J. (2019) Prevalence and diversity of antibiotic resistance genes in Swedish aquatic environments impacted by household and hospital wastewater. Front Microbiol 10: 688.

Kumpitsch, C., Koskinen, K., Schöpf, V., and Moissl-Eichinger, C. (2019) The microbiome of the upper respiratory tract in health and disease. BMC Biol 17: 1– 20.

Li, J., Cao, J., Zhu, Y., Chen, Q., Shen, F., Wu, Y., et al. (2018) Global Survey of Antibiotic Resistance Genes in Air. Environ Sci Technol 52: 10975–10984.

Li, Z., Xia, J., Jiang, L., Tan, Y., An, Y., Zhu, X., et al. (2021) Characterization of the human skin resistome and identification of two microbiota cutotypes. Microbiome 9: 47.

Lira, F., Vaz-Moreira, I., Tamames, J., Manaia, C.M., and Martínez, J.L. (2020) Metagenomic analysis of an urban resistome before and after wastewater treatment. Sci Reports 2020 101 10: 1–9.

Maestre-Carballa, L., Lluesma Gomez, M., Angla Navarro, A., Garcia-Heredia, I., Martinez-Hernandez, F., and Martinez-Garcia, M. (2019) Insights into the antibiotic resistance dissemination in a wastewater effluent microbiome: bacteria, viruses and vesicles matter. Environ Microbiol.

Methé, B.A., Nelson, K.E., Pop, M., Creasy, H.H., Giglio, M.G., Huttenhower, C., et al. (2012) A framework for human microbiome research. Nature 486: 215–221.

Naidoo, Y., Valverde, A., Cason, E.D., Pierneef, R.E., and Cowan, D.A. (2020) A clinically important, plasmid-borne antibiotic resistance gene (β-lactamase TEM-116) present in desert soils. Sci Total Environ 719: 137497.

Nayfach, S. and Pollard, K.S. (2015) Average genome size estimation improves comparative metagenomics and sheds light on the functional ecology of the human microbiome. Genome Biol 16: 1–18.

Nelson, R.E., Hatfield, K.M., Wolford, H., Samore, M.H., Scott, R.D., Reddy, S.C., et al. (2021) National Estimates of Healthcare Costs Associated With Multidrug-Resistant Bacterial Infections Among Hospitalized Patients in the United States. Clin Infect Dis 72: S17–S26.

O’Neill, J. (2016) Tackling drug-resistant infections globally: final report and recommendations, United Kingdom.

Olivares, J., Bernardini, A., Garcia-Leon, G., Corona, F., Sanchez, M.B., and Martinez, J.L. (2013) The intrinsic resistome of bacterial pathogens. Front Microbiol 4: 103.

Palleja, A., Mikkelsen, K.H., Forslund, S.K., Kashani, A., Allin, K.H., Nielsen, T., et al. (2018) Recovery of gut microbiota of healthy adults following antibiotic exposure. Nat Microbiol 3: 1255–1265.

R Core Team (2014) R: A Language and Environment for Statistical Computing. R Found Stat Comput.

Reygaert, W.C. (2018) An overview of the antimicrobial resistance mechanisms of bacteria. AIMS Microbiol 4: 482–501.

Rodriguez-Mozaz, S., Vaz-Moreira, I., Varela Della Giustina, S., Llorca, M., Barceló, D., Schubert, S., et al. (2020) Antibiotic residues in final effluents of European wastewater treatment plants and their impact on the aquatic environment. Environ Int 140: 105733.

Rowe, W.P.M., Baker-Austin, C., Verner-Jeffreys, D.W., Ryan, J.J., Micallef, C., Maskell, D.J., and Pearce, G.P. (2017) Overexpression of antibiotic resistance genes in hospital effluents over time. J Antimicrob Chemother 72: 1617–1623.

Salyers, A.A., Gupta, A., and Wang, Y. (2004) Human intestinal bacteria as reservoirs for antibiotic resistance genes. Trends Microbiol 12: 412–416.

Shaw, W. V., Packman, L.C., Burleigh, B.D., Dell, A., Morris, H.R., and Hartley, B.S. (1979) Primary structure of a chloramphenicol acetyltransferase specified by R plasmids [31]. Nature 282: 870–872.

Ventola, C.L. (2015) The antibiotic resistance crisis: causes and threats. P T J 40: 277– 83.

Wickham, H. (2016) ggplot2: Elegant Graphics for Data Analysis, Springer-Verlag New York.

Wright, G.D. (2010) Q&A: Antibiotic resistance: where does it come from and what can we do about it? BMC Biol 2010 81 8: 1–6.

Zhang, T., Li, X., Wang, M., Chen, H., Yang, Y., Chen, Q. lin, and Yao, M. (2019) Time-resolved spread of antibiotic resistance genes in highly polluted air. Environ Int 127: 333–339.

